# REPOP: bacterial population quantification from plate counts

**DOI:** 10.1101/2025.04.01.644179

**Authors:** Pedro Pessoa, Carol Yunxiao Lu, Stanimir Asenov Tashev, Rory Kruithoff, Douglas P. Shepherd, Steve Pressé

## Abstract

Bacterial counts from native environments, such as soil or the animal gut, often show substantial variability across replicate samples. This heterogeneity is typically attributed to genetic or environmental factors. A common approach to estimating bacterial populations involves successive dilution and plating, followed by multiplying colony counts by dilution factors. This method, however, overestimates the heterogeneity in bacterial population because it conflates the inherent uncertainty in drawing a subsample from the total population with the uncertainty in the sample arising from biological origins. In other words, this approach may obscure features that may otherwise be present in the data hinting at the presence of genuine subpopulations. For example, in plate counting applied to *C. elegans* gut microbiota, observed multimodality is often interpreted as large host-to-host variance, while the randomness introduced by measurement is frequently ignored. To explicitly account for the uncertainty introduced by dilution and plating randomness, we introduce REPOP, a PyTorch-based library to REconstruct POpulations from Plates within a Bayesian framework. Beyond simple cases, REPOP addresses more complex scenarios, including multimodal populations and correcting the mathematically subtle, but experimentally relevant, bias introduced by excluding plates deemed too crowded to distinguish individual colonies. We demonstrate REPOP’s ability to resolve distinct population peaks otherwise obscured by standard multiplication methods. Applications to both simulated and experimental datasets, including bacterial samples of different concentrations and ones from the gut microbiota of *C. elegans*, show that REPOP accurately recovers the underlying multimodality by properly accounting for error propagation, where naive multiplication fails.

REPOP is available on GitHub: https://github.com/LabPresse/REPOP.

## Introduction

Plate counting is fundamental to microbiology in estimating the number of living microorganisms in a sample (***Corry et al., 2011; Smith and Brown, 2021***). For example, it is essential in food safety testing (***Vial et al., 2019***), water quality assessments (***Bartram et al., 2003; Some et al., 2021***), amongst many other applications (***Wang et al., 2020; Tamargo et al., 2022; Martini et al., 2024; Taylor et al., 2022***) that require flexible quantification of microbial loads.

Briefly, plate counting involves serially diluting an original sample to ensure an appropriate colony density, allowing for the accurate enumeration of colonies for each sample (as summarized in Fig. 1a). Subsequently, the diluted samples are spread over nutrient-rich agar plates and incubated. After incubation, each viable organism (typically a bacterium) spread over the plate forms a colony, or in some cases a plaque (***Marine et al., 2020***). All of the colonies or plaques on the plate are then enumerated as colony forming units (CFUs) or plaque forming units (PFUs), respectively. The degree of dilution (represented by the dilution factor) is typically set in such a way as to collect a statistically precise number of colonies but simultaneously avoiding colony overlap – usually aiming for 30 to 300 CFUs in a plate with a diameter of 10 cm. However, when studying a poorly characterized or highly variable bacterial population, collecting plates that are overcrowded is inevitable. Challenges related to counting crowded plates were only recently mitigated by AI-inspired tools (***Geissmann, 2013; Khan et al., 2018; Albaradei et al., 2020; Zhang et al., 2021***).

**Figure 1.**
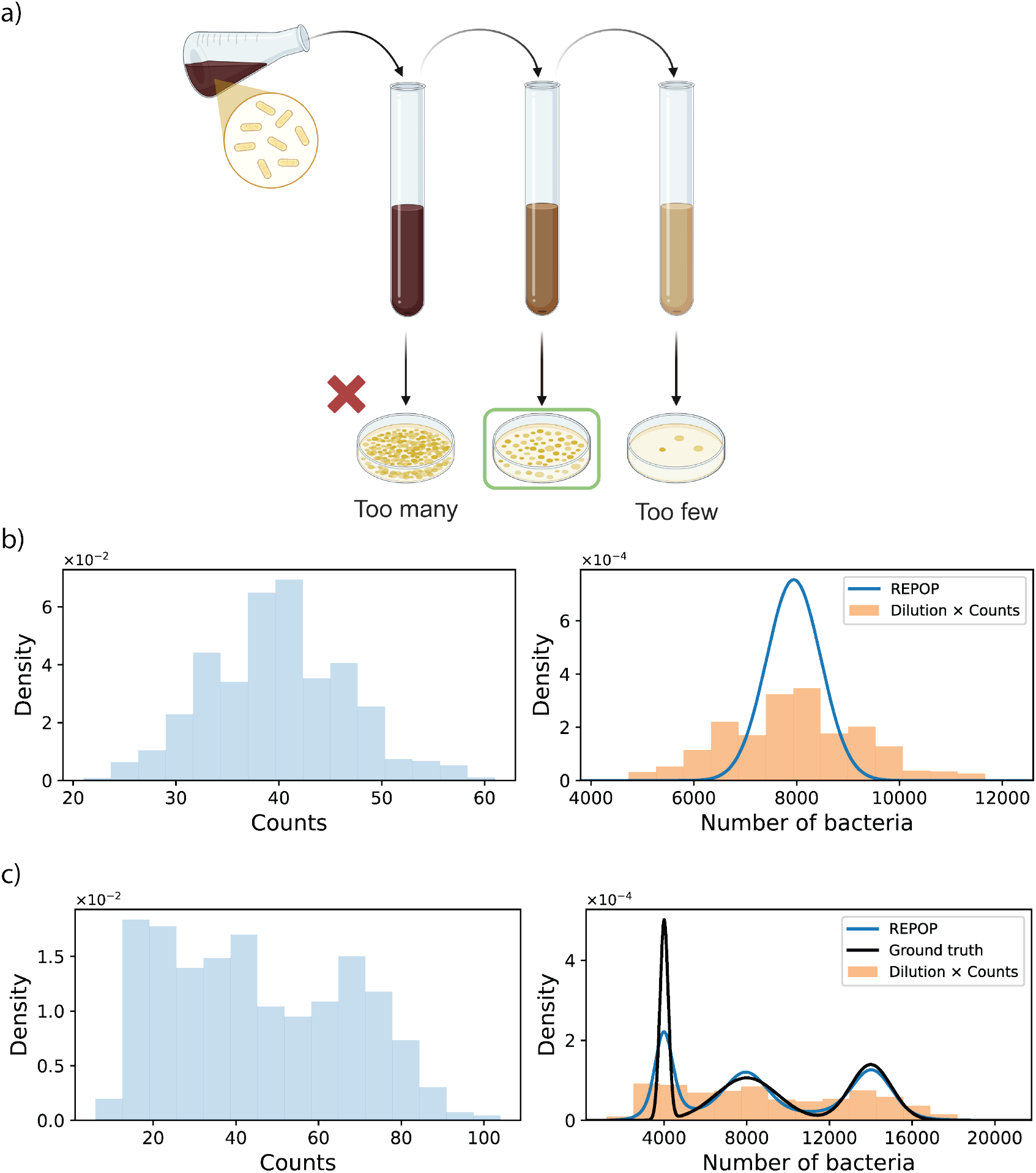
Visual description of the dilution and plate counting process and the need for its analysis. a) A sample of volume *V* is mixed with sterile medium in a larger vial (*e*.*g*., 19*V*, for a total dilution of 20). A portion of volume *V* of this diluted sample is then spread on a plate and incubated. Further tenfold dilutions are performed by transferring *V* into a vial with 9*V* of medium before plating. The process continues until a reasonable number of counts is obtained. b) The histogram on the left shows simulated counts across 1000 plates, obtained by diluting samples from the ground truth distribution by a factor of 200. The usual method of multiplying the counts by the total dilution factor produces the orange histogram (in the right), which is significantly broader than the ground truth (black line, right) due to additional stochasticity introduced by dilution and plating. The ground truth distribution is a Gaussian with a mean of 8000 and a standard deviation of 500. REPOP’s reconstruction (blue) corrects the broadening. c) A similar simulation to (b) where the ground truth population distribution is trimodal (parameters in Fig. 3). Here, the stochasticity introduced by dilution and plating obscures the three peaks, making them difficult to distinguish in the observed counts. Nevertheless, REPOP successfully reconstructs the trimodal distribution.

Beyond simple quantification, plate counting is often used to assess population variation within a single microbial species, the degree of phenotypic variability (***Martino et al., 2016; De Martino, 2017; Pecht et al., 2019; Muntoni et al., 2022***) and environmental resource availability (***Armitage and Jones, 2019; Blanchet et al., 2020***). This stems from interpreting multimodality – several samples exhibiting high and several others low counts – or general heterogeneity in plate counts as reflecting population differences in colony formation, growth rates, or stochastic expression of traits (***Veening et al., 2008; Moore et al., 2013***). This heterogeneity can then be used to understand ecological dynamics of bacterial populations (***Mao and Lu, 2016; Armitage and Jones, 2019; Werner et al., 2022; Goberna and Verdú, 2022; Pinto et al., 2022***) such as the competition of populations inside the gut of *Caenorhabditis elegans* (*C*.*elegans*) (***Lu et al., 2026; Vega and Gore, 2017***).

On account of this, learning the variability from sample to sample is critical in accurately capturing a population’s heterogeneity. Thus, in order to draw biological conclusions, one must disentangle the effects of the stochasticity involved in the measurement (e.g., the randomness involved in the process of dilution and plating) from the heterogeneity in the original sample’s population arising from the underlying biological or ecological variation.

Naively, in dilution and plating, the total number of viable bacteria in the original sample is usually estimated by multiplying the number of colonies by the total dilution factor for each plate, resulting in a histogram of the population in the original sample. This process significantly overestimates the true variability in the bacterial count within the samples and can even obscure the multimodality in reconstructed population count histograms, as demonstrated in Fig. 1b and c.

Indeed, this higher variance introduced in quantification is a major disadvantage of plate counting when compared to other methods, such as those based on flow cytometry or DNA quantification (***Van Nevel et al., 2017; Wilkinson, 2018; Hansen et al., 2020; Weitzel et al., 2021; Tracey et al., 2023; Sun et al., 2024***), which have the ability to count cells without dilution. Yet, plate counting remains the gold standard in several areas of microbiological research due to its ability to quantify only viable cells (***Davey and Guyot, 2020; International Organization for Standardization, 2013; Bartram et al., 2003***). In contrast, flow cytometry can only distinguish viable cells under specific conditions. It typically relies on fluorophore-based methods, assuming that a high intracellular concentration of non-membrane-permeable fluorescent probes indicates a compromised membrane and, therefore, a lack of viability (***Taimur et al., 2020; Davey and Guyot, 2020***). These challenges underscore why plate counting remains relevant, and improvements in the technique have wide downstream implications.

To overcome the limitations of traditional plate count estimation, we developed REPOP (REconstruct POpulations from Plates), a software package that employs a Bayesian approach that rigorously accounts for the stochasticity introduced by dilution and plating. By modeling these statistical processes, REPOP infers both the full probability distribution of the original samples, preserving their inherent heterogeneity, as demonstrated in Fig. 1b. Notably, REPOP can detect the multimodal structure of the data even when such multimodality is not readily apparent from dilution count histograms, as shown in Fig. 1c. REPOP is developed in Python and built on PyTorch (***Paszke et al., 2019***). Although PyTorch was originally developed for neural networks, it is also well suited for the type of optimization and GPU-accelerated calculations required here. REPOP is pip-installable and can take advantage of a GPU when available.

REPOP is available on GitHub (***Pessoa, 2025***), where users can find recipe-like tutorials, including examples of how to reconstruct the underlying population distribution and obtain quantities such as quantiles. A schematic overview of the workflow is shown in Fig. 2. Briefly, the input data consist of a sequence of observed colony counts together with their respective dilution factors, where each pair corresponds to a single biological sample. These data are passed to the inference pipeline, which evaluates the likelihood and reconstructs the distribution of the original bacterial population across samples. The main output is the inferred population distribution, along with summary statistics such as moments or quantiles.

**Figure 2.**
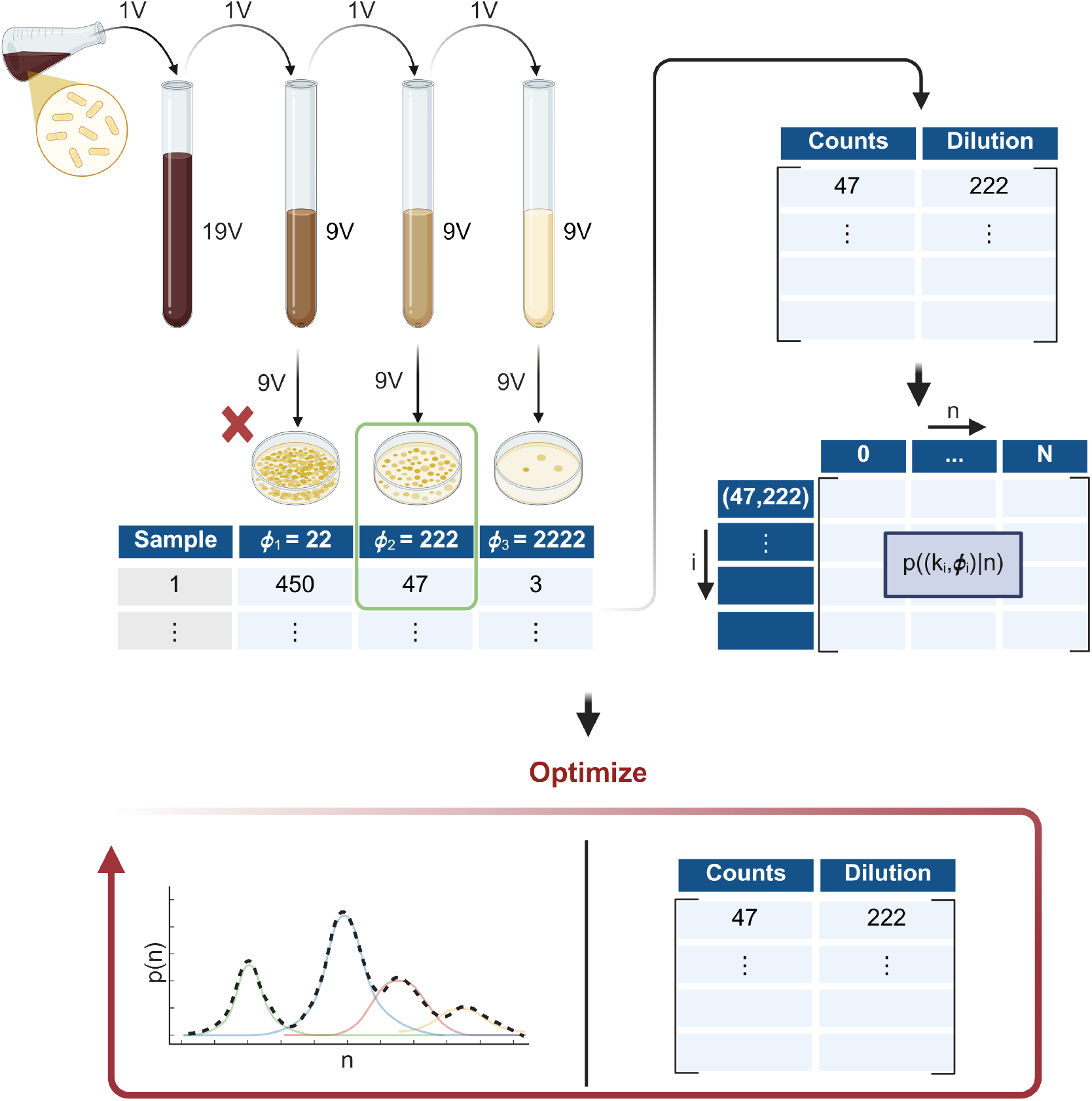
REPOP workflow from plate counts to reconstructed population statistics. The input consists of observed colony counts and their corresponding dilution factors, organized as pairs (*k*_*i*_, *ϕ*_*i*_), each arising from a single biological sample. From these data, REPOP evaluates the probability of each observed dilution count pair conditioned on the underlying bacterial number, *p*(*k*_*i*_, *ϕ*_*i*_ ∣ *n*_*i*_), thereby capturing the stochasticity introduced by dilution and plating. These likelihood contributions are then combined to reconstruct the distribution of the original bacterial population across samples.

When presenting our method, we address a mathematically subtle point that may qualitatively affect the inferred counts: the cutoff for “uncountable” on a plate. That is to say, when multiple dilutions are performed in experiments to obtain plates with a large quantity of CFUs, plates that exceed a preset colony cutoff (a number beyond which colonies are considered too numerous to count) are excluded. While this practice is common, failing to properly account for the exclusion of such plates can introduce bias in the inferred population distribution.

Here, we will explain how REPOP is able to integrate previously ignored over-populated plates exceeding the cutoff in a more sophisticated albeit requisite iteration of our simpler method.

Why requisite? Under the very general assumption that there is a nonzero probability of the population sizes being larger than the cutoff, no matter how low we set our dilution factor, a plate may still probabilistically exceed the countable cutoff, though increasingly rarely. Thus, if we can deal with over-populated plates, then we may collect better statistics on colony counts by avoiding too large a dilution.

To demonstrate REPOP’s applicability to real world datasets, we analyzed two distinct experimental systems. First, we applied REPOP to plate counting data obtained from controlled mixtures of *Escherichia coli* (*E. coli*) populations at different optical densities. Without being provided the optical density condition from which each individual sample was drawn from, REPOP successfully reconstructed the mixture distribution of bacterial abundances across the different conditions. Second, we employed REPOP to study bacterial colonization dynamics in individual *C. elegans* hosts. By analyzing plate counts of gut bacterial populations over several days post feeding, REPOP revealed the progressive increase in bacterial load and captured the multimodal structure associated with host heterogeneity. These results highlight REPOP’s ability to disentangle true biological variability from stochastic sampling noise in complex, real experimental contexts.

## Methods

### Statistical description of plate counting

We first describe the process of serial dilutions in mathematical terms. From Fig. 1a, we observe that, on each plate, a fraction of the volume of the initial sample, equal to the inverse of the dilution factor (*ϕ*), is spread over the plate.

In practice, a sample is first collected from the biological system and then mixed as part of the dilution procedure prior to plating, resulting in a well-mixed solution from which a subvolume corresponding to a fraction ^1^/_*ϕ*_ of the total volume is transferred to the plate. Consequently, each individual bacterium in the diluted sample has probability ^1^/_*ϕ*_ of being transferred onto the plate.

We denote the total number of bacteria in the original sample as *n*. Assuming each bacterium is independently and equally likely to be transferred to the final plate and form a colony, the resulting number of colonies, denoted as *k*, follows a binomial distribution with parameters *n* (the number of trials) and 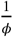(the probability of success in each trial), expressed as

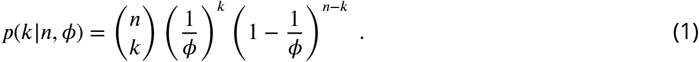

Here, for the numbers usually used in plate counting, the stochasticity introduced by the dilution is far from negligible. From (1), the variance of *k* is given by 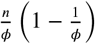, while the mean is 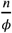. Thus, for large dilution factors, the variance is approximately equal to the mean. In practice, we expect colony counts to remain below 300. For example, if the expected count is 150, then the binomial variance is approximately 150, corresponding to a standard deviation of 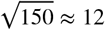. Therefore, the observed count *k* will typically lie within one standard deviation of the mean, approximately in the interval [138, 162]. This corresponds to a relative error of approximately 8%. Similarly, for an expected count of 50 (smaller but still reasonable) the standard deviation of *k* is approximately 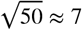. Therefore, for a case where the expected number is 50, one standard deviation corresponds roughly to the interval [43, 57], which gives a relative error of approximately 14%.

### Bayesian inference

For now, we assume that each sample (drawn from the sample at the previous dilution) gives rise to one plate. The dataset obtained consists of the sequence of plate counts, *k*_*i*_ and their respective dilution factors *ϕ*_*i*_ for each replicate *i*. We denote the complete dataset as 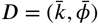 with 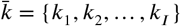 and 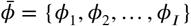, with each pair (*k*_*i*_, *ϕ*_*i*_) generated independently as they arise from different samples.

Our overall goal is to learn the probability distribution for the number of bacteria in the original samples, *n*, from the dataset *D*. To achieve this, we assume *a priori* that the distribution over *n* follows a model of population distribution with a set of parameters *θ*, written as *p*(*n* |*θ*).

Later, we will consider both unimodal and multimodal population distributions, *p*(*n*| *θ*). First, assuming that the population distribution is unimodal, we represent *p*(*n*|*θ*) as a simple Gaussian distribution, where *θ* reflects the mean and standard deviation. Second, in order to model a multimodal distribution, we invoke non-parametric models which provide a means to describe multimodal probability distributions for which the number of modes is unknown *a priori* (***Pressé et al., 2010; Jazani et al., 2019; Tavakoli et al., 2020; Kilic et al., 2021; Sgouralis et al., 2024; Pressé and Sgouralis, 2023***). More specifically, we assume a non-parametric Gaussian mixture, where *θ* represents a (potentially infinite) set of means, standard deviations, and mixture weights. Importantly, in large datasets, non-parametric models can flexibly adapt to approximate probabilities that do not strictly follow a predefined functional form (***Bryan IV et al., 2020; Pressé and Sgouralis, 2023; Pessoa et al., 2025***).

The goal of reconstructing the distribution of bacteria in the original samples then becomes one of learning the underlying parameters (*θ*) from plate count data (*D*). In other words, we want to learn the probability of *θ* conditioned on *D, p*(*θ*|*D*). This probability is called a posterior. As we will see later, calculating the posterior will require the use of *p*(*n* | *θ*), which is uni- or multimodal, and the use of *p*(*ϕ*_*i*_ | *n*) when cutoffs on countability of plates are used in order to eliminate plates containing too many colonies.

REPOP is applied here to three cases of increasing complexity: the first (Model 1) is the simplest, assumes unimodal *p*(*n* | *θ*) without cutoffs. In Model 2, we keep no cutoff in counts, but generalize *p*(*n* | *θ*) to a multimodal (which simplifies to the unimodal Model when all but one component is considered). In Model 3, we keep the most general multimodal distribution for *p*(*n* |*θ*), but expand the counting method to account for the cutoff, resulting in some plates not being counted due to exceeding the cutoff. We summarize the differences among the three mechanisms in Table 1.

**Table 1.**
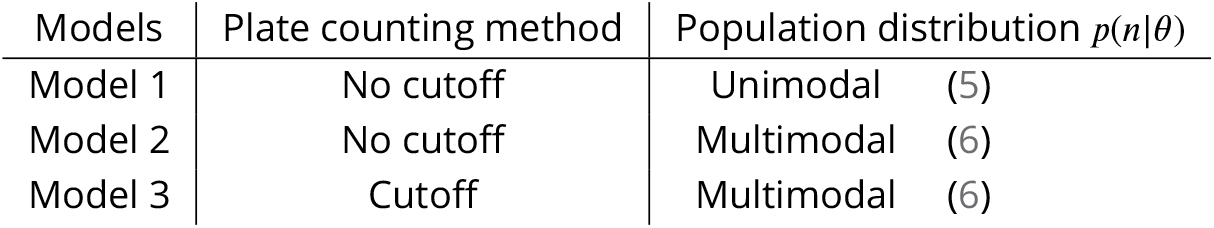
For ease of reference, we summarize the differences among the three models taken into account in the present article.

For all models highlighted above, in order to compute *p*(*θ* |*D*), we need the likelihood, *p*(*D*| *θ*), then related to *p*(*θ*|*D*) through Bayes’ theorem

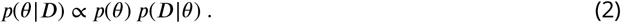

Here *p*(*θ*) is the prior. Further details on the prior are given in the Appendix 1. Once we calculate probability *p*(*θ* | *D*), we may maximize this probability to obtain the maximum *a posteriori* (MAP) value, *θ*^MAP^ (as we describe later).

### Population versus measurement stochasticity

In the following sections, we describe how the likelihood, *p*(*D* ∣ *θ*), is computed. We denote the dataset as 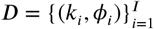, where each pair corresponds to the colony count and associated dilution factor for sample *i*. Assuming samples are uncorrelated, except in sharing the same underlying ecology, they may be treated as independent and identically distributed, so that the likelihood factorizes as

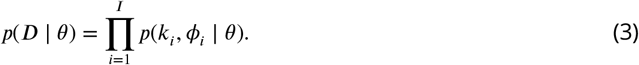

However, each observed plate count arises from an underlying (and unobserved) number of bacteria in the original sample, which we denote by *n*_*i*_. This latent variable captures the true population size for sample *i* and is the quantity directly informed by the ecological variability. Thus, we can rewrite each term in the likelihood by marginalizing over *n*_*i*_,

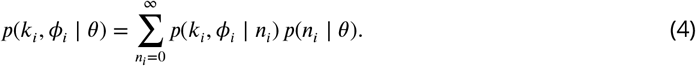

This decomposition makes explicit the two sources of stochasticity. The term *p*(*n*_*i*_ ∣ *θ*) describes the population distribution, encoding the intrinsic variability across samples, while *p*(*k*_*i*_, *ϕ*_*i*_ ∣ *n*_*i*_) describes the measurement process, capturing the stochastic effects of dilution and plating. The following two subsections explain these two ingredients in detail. In particular, the population model *p*(*n*_*i*_ ∣ *θ*) is flexible and must be chosen to reflect the underlying biological or ecological structure of interest, whereas the measurement model *p*(*k*_*i*_, *ϕ*_*i*_ ∣ *n*_*i*_), explained in the following subsection, is dictated by the experimental protocol itself.

In the following sections, we introduce a hierarchy of models for *p*(*n*_*i*_ ∣ *θ*), ranging from simple unimodal distributions to flexible non-parametric multimodal representations. While REPOP provides general purpose choices for these models, users interested in specific ecological hypotheses may substitute alternative forms for *p*(*n*_*i*_ ∣ *θ*) without modifying the measurement model. That is, REPOP can also be used to extract only the measurement component, *p*(*k*_*i*_, *ϕ*_*i*_ ∣ *n*_*i*_), whose implementation is described in detail on GitHub (***Pessoa, 2025***).

### Models for the population distribution

#### Unimodal distribution (Model 1)

For Model 1, we can mathematically represent the probability of the true population count *n*_*i*_ given the distribution parameters *θ* as:

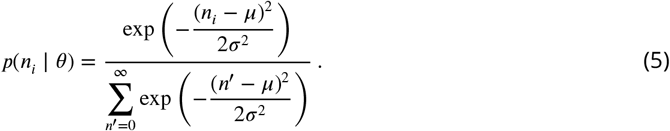

Here, *θ* includes the mean, µ, and standard deviation, σ, of the integer Gaussian *p*(*n*_*i*_| *θ*), *θ* = {µ, σ}.

The denominator above is necessary because we are dealing with (positive) integer Gaussians and will approach the usual 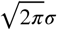 when *μ* – 5σ≫1. For computational reasons, we assume that the probability of having a number of bacteria, *n*, larger than twice the largest number found by multiplying CFU counts per dilution is approximately zero. Thus, the infinite upper bound over the sum in the denominator of (5) is substituted for *N*^∗^ = 2 max(*k*_*i*_*ϕ*_*i*_). Note that, by using the integer Gaussian in (5), we truncate the negative support of a Gaussian distribution, meaning we assume that even if we have a pair of *µ* and *σ* that assigns non-negligible probability to counts *n*_*i*_ ≤ 0, those probabilities are set to zero, and the distribution is renormalized accordingly.

#### Multimodal distribution non-parametric inference (Models 2 and 3)

Naturally, the choice of modeling *n* through a single Gaussian component, Model 1, already commits us to assuming a unimodal distribution. However, if bacteria are present across replicates in a multimodal distribution (such as observed in some studies of *C*.*elegans* gut microbiota (***Vega and Gore, 2017; Taylor and Vega, 2024; Martini et al., 2024***), where some replicates have few bacteria and others have more based on rare colonization events ***Vega and Gore*** (***2017***); ***Taylor and Vega*** (***2024***)) how many modes should we consider?

This consideration immediately forces upon us an important motivation for considering a much more general choice for *p*(*n*| *θ*), namely a non-parametric Gaussian mixture. Loosely speaking, non-parametric Gaussian mixture models do not assume a fixed number of parameters (or components) beforehand. Instead, they adapt and determine the appropriate number of components as warranted by the data. Knowledge of non-parametric mixture models, while subtle and detailed in ***Pressé and Sgouralis*** (***2023***) and references therein, is not required for running REPOP. In such a model, *θ* encompasses the means 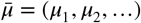, standard deviations 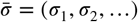, and mixture weights (the relative weights of each Gaussian component with the weights summing to unity) 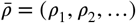, such that 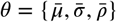. In this non-parametric model, the probability of having a sample of *n* bacteria is given as

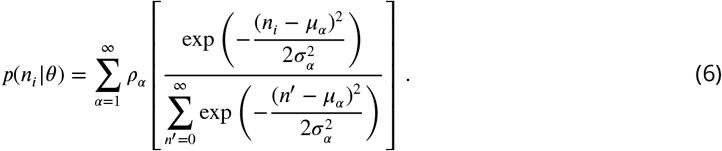

When performing non-parametric inference, the choice of prior *p*(*θ*) is key. A well-chosen prior can add computational efficiency and avoid runaway pathologies associated with assigning every datapoint a different mixture component. A common choice of prior for the infinite mixture model satisfying both of these criteria is the Dirichlet process prior (***Pressé and Sgouralis, 2023***); see Appendix 1 for more details. While a finite truncation in the number of components can be avoided (***Walker, 2007; Kalli et al., 2009***), for computational reasons, here we use a finite truncation (defaulted to the smallest value among 25 and the square root of the total number of samples, though modifiable, in REPOP) for the sum over the components, α, in (6).

### Calculating the likelihood

Here we explain how to calculate the likelihood, *p*(*D* | *θ*), appearing in (2) and start with Models 1 and 2 and move to Model 3, where the cutoff is taken into consideration.

#### Likelihood for Models 1 and 2

In the Models where cutoffs are not taken into consideration (*i*.*e*., where it is assumed that the colonies on a plate can always be counted and plates are never rejected for having too many colonies), the calculation of the total likelihood for all counted plates from the original number of bacteria in one sample, *n*, is given by (1). However, *n*_*i*_ is itself a realization of the population distribution *p*(*n*|*θ*). Thus, the probability that a plate with *k*_*i*_ counts was generated from a (fixed) dilution *ϕ*_*i*_ is given by

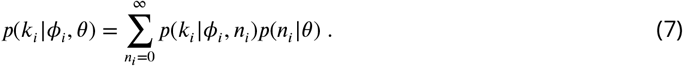

Here, the dilution factor is treated as fixed rather than as a random variable because it is assumed that the colonies can always be counted. Within the summation in (7), each factor *p*(*k*_*i*_ *|*ϕ_*i*_, *n*_*i*_) within the summation given by (1) and *p*(*n*_*i*_ |*θ*) given by the respective population model as presented in Models 1–3 in the previous subsection.

This allows us to write Bayes’ theorem (2) as

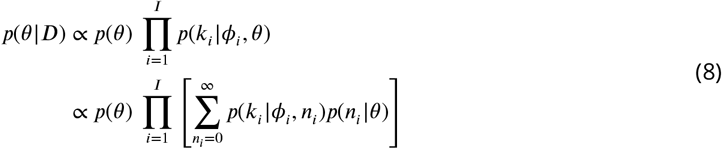

where *I* is defined as the total number of replicates. Note that while this summation is truncated to twice the naively reconstructed maximum from observed CFU counts 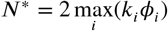, these sums can still remain computationally intensive. Thus, while REPOP can run on CPUs, we automatically leverage GPU acceleration to enable faster evaluation of these summations if a GPU is available.

#### Likelihood for Model 3

For Model 3, we assume that the number of colonies may exceed a certain cutoff, *k*_CO_, and that such plates are not counted. Typically, for each sample, only one of the plates generated in the process described in Fig. 1a is reported. In other words, for each collected sample only the counts (and dilution) of one of the plates at the first dilution resulting in a countable number of colonies (below a pre-defined cutoff) is recorded as a datapoint. This is illustrated in Fig. 1a.

This process is different from the previous models. If a dilution different from the first dilution in the dilution schedule is reported, this implies that all lower dilutions therefore resulted in plates whose colony count (a stochastic variable) already exceeded the cutoff *k*_CO_.

To obtain the posterior over *θ*, it is necessary to know the dilution schedule from the plating experiment. We refer to the full set of these as Φ = {*ϕ*^1^, *ϕ*^2^, …, *ϕ*^*M*^ } ordered from lowest to highest dilution factor. In the example given in Fig. 1a, we have *ϕ*^1^ = 22, *ϕ*^2^ = 222, and *ϕ*^3^ = 2222. While, mathematically, we could extend this indefinitely by sequentially plating dilutions that are factors of 10 larger, in practice only a finite number of plates can be produced, thus justifying the upper limit *M*. It is expected that the dilution schedule leads to dilution values sufficiently large that the approximate number of colonies recovered approaches zero. Expanding upon the previous models, it is now important to report not only the counts but also the associated plate index. The datapoint is now reported as (*k*_*i*_, *m*_*i*_), which means that for the *i*-th sample, the plate diluted by a total factor of 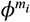had *k*_*i*_ counts and all plates with smaller dilution had counts larger than the cutoff *k*_CO_.

With the set of all possible dilutions Φ explicitly considered, we incorporate this dependence into the calculation of all relevant probabilities. Thus, the total posterior for the full dataset is given by

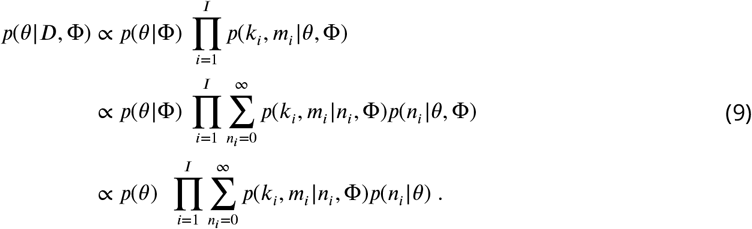

Here, we used the fact that the population distribution and its parameters, *θ*, are independent of the dilutions selected to plate, Φ. Therefore, *p*(*θ*| Φ) = *p*(*θ*) is the original prior and *p*(*n*_*i*_ | *θ*, Φ) = *p*(*n*_*i*_ | *θ*) is given by the respective population model. The remainder of this subsection focuses on how to calculate the likelihood of each datapoint, *p*(*k*_*i*_, *m*_*i*_ | *n*_*i*_, Φ), within the summation above. To obtain this likelihood, we revisit the dilution and plating process. As described in Fig. 1a, a plate is generated for each dilution within the set of dilutions Φ. For each sample, labeled *i*, the number of colonies on the plate with dilution *ϕ*^*m*^ is denoted as 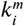.

Since all plates originate from the same original sample (with a finite number of bacteria, *n*_*i*_), the values of 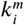 for the same *i* are, in principle, not independent. Instead, the set of all 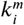 is jointly sampled from a single multinomial distribution. However, as each bacterium has an independent probability of 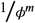 of appearing on the *m*-th plate, the correlations between the multiple 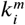 arise solely due to the finite number of total bacteria in the sample. Consequently, as long as the fraction of plated bacteria remains significantly smaller than the total sample size (*i*.*e*.,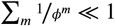, the number of plated bacteria can be treated as independent binomial random variables.

Under this assumption, each count 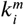 can be modeled as a binomial realization of the original number of bacteria *n*_*i*_ with dilution factor *ϕ*^*m*^, as described in Eq. (1). Thus, the likelihood of observing 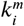 is written as:

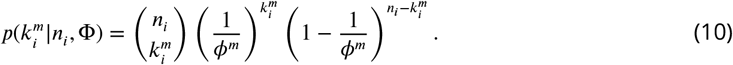

As previously described, the datapoint (*k*_*i*_, *m*_*i*_) corresponds to the counts and the index of the first value within the sequence 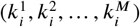 that satisfies the condition 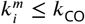, where *k*_CO_ is the cutoff. To formalize this, we define *m*_*i*_ as the (superscript) index of the first plate where the counts fall within the cutoff:

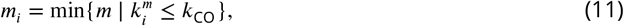

from which the observed datapoint (*k*_*i*_, *ϕ*_*i*_) is obtained with:

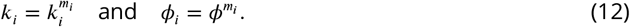

To avoid confusion, hereinafter we express the likelihood in terms of the plate index *m*_*i*_ rather than the corresponding dilution factor *ϕ*_*i*_, since the ordered dilution schedule Φ establishes a one-to-one correspondence between them.

Within this convention. the likelihood, *p*(*k*_*i*_, *m*_*i*_ | *n*_*i*_, Φ), is written as

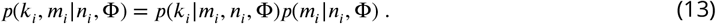

Once more, we proceed by showing how to calculate each factor within the product above. Starting from the last factor, *p*(*m*_*i*_ | *n*_*i*_, Φ), we note that if the plate with index *m*_*i*_ is reported, we must have that both 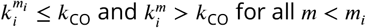,

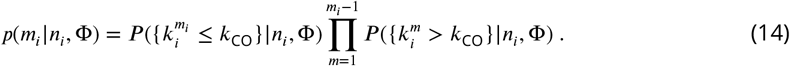

Here, we use the capitalized *P* to represent that we are calculating the probability that a variable is within a set, rather than the probability of a single possible value. In this specific model, what we are asking is the probability that 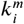 is smaller than or equal to *k*_CO_. This is given by summing the probability of every value of 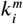up to *k*_CO_, meaning

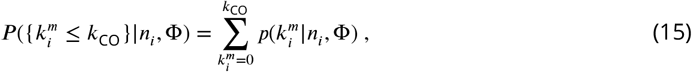

so (14) becomes

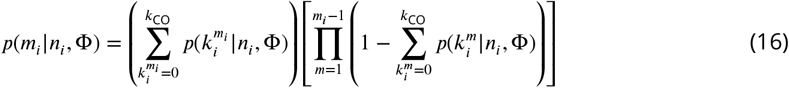

with each 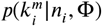 given by (10).

Then, for the remaining factor in (13), *p*(*k*_*i*_ |*m*_*i*_, *n*_*i*_, Φ), since counts above the cutoff are not reported as data, the likelihood of finding *k*_*i*_ colonies in a plate diluted by a total factor 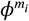 is re-scaled to account for the fact it was truncated at the cutoff

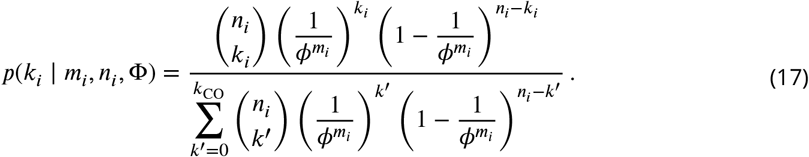

for *k*_*i*_ ≤ *k*_CO_ and zero otherwise.

Thus, in order to obtain the posterior (9), we need to substitute the likelihood per datapoint, *p*(*k*_*i*_, *m*_*i*_|*n*_*i*_, Φ) in (13), which is computed using (17) and (16).

### Bayesian MAP

In Bayesian inference, we compute the posterior distribution *p*(*θ*|*D*) as it captures the updated probability over the model parameters after observing the data. However, posteriors are often complex and high-dimensional (technically infinite in non-parametric models), making direct computation and visualization challenging. To address this, we often resort to sampling methods to approximate the posterior. However, these samplers can also require long computation.

Since our objective here is to create a tool that is quick and easy to run, we resort to maximum *a posteriori* (MAP) estimation. This reduces the problem of knowing the distribution over *θ* to identifying the optimal *θ*. In this model, we maximize *p*(*θ*|*D*) and denote the maximum value as *θ*^MAP^,

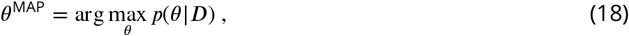

with *p*(θ|*D*) given by (8) for Models 1 and 2, or (9) for Model 3. For our purposes here, we say that *p*(*n*|θ^MAP^) is the distribution of *n* we learned from the data. In REPOP, the maximum value θ^MAP^ is found using optimization libraries within PyTorch (***Paszke et al., 2019***).

## Results

In the previous section, we have constructed the posterior *p*(*θ* | *D*) across unimodal and multimodal population distributions, and by using a likelihood that either assumes all plates are countable (Model 1-2) or considers plates exceeding the cutoff (Model 3).

### When all plates are considered countable

For the first model, we generate simulated data drawn from a single Gaussian with equal dilution factors across all samples. This assumes that the dilution factor used (200) is able to generate a countable number of colonies in all samples. We also assume a unimodal Gaussian, as in Model 1. The reconstruction obtained by REPOP was shown earlier in Fig. 1b, where we see that REPOP corrects for the overestimation of variance introduced by the stochasticity in dilution and plating.

In the more complex multimodal Gaussian case (Model 2), we further observe that REPOP uncovers the true underlying distribution’s peaks with greater resolution and sensitivity to multimodality than by using naive multiplication. As an example, in Fig. 3 we generate simulated data for a mixture of Gaussians with three modes.

**Figure 3.**
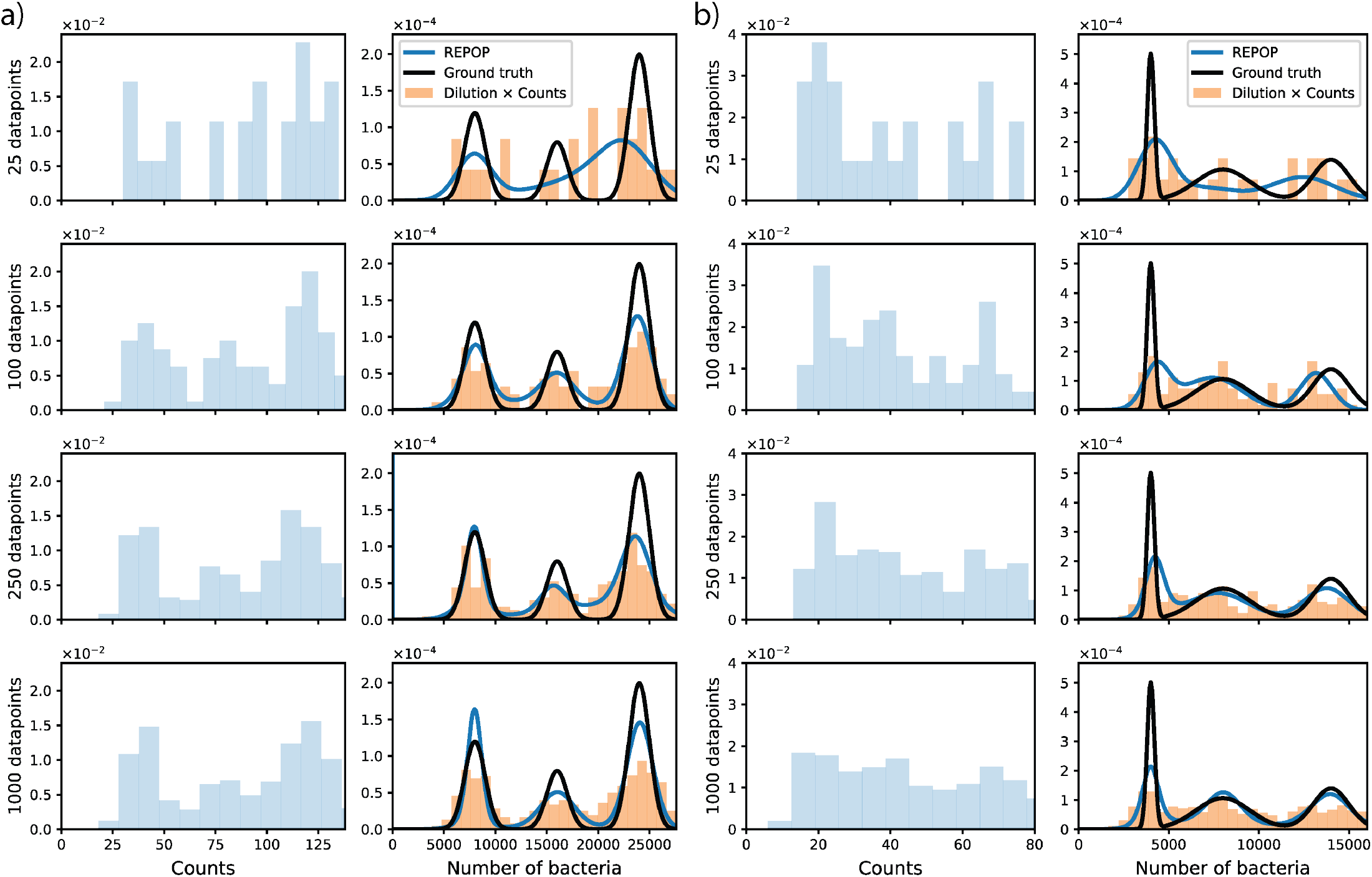
REPOP can reconstruct multimodal populations even when they are not directly observable by naively multiplying colony counts by dilution. Similar to Fig. 1b and c, we show above the observed colony counts, the naïve estimate obtained by multiplying counts by the dilution factor, and the reconstruction produced by REPOP, compared to the ground truth. In both cases, the population size *n* is drawn from a mixture of three Gaussians, shown as the black curves. In a), the mixture has means (4000, 8000, 14000), standard deviations (200, 1500, 1000), and weights (0.25, 0.4, 0.35). In b), the mixture has means (8000, 16000, 24000), standard deviations (1000, 1000, 1000), and weights (0.3, 0.2, 0.5). Observed counts are sampled from (1), with dilution factor 200, and are shown as histograms on the left. In (a), a trimodal structure is visible in the observed count histogram. However, in b), the trimodal structure is strongly obscured by stochasticity from dilution and plating, making it difficult to discern even in large datasets. Despite this, applying REPOP to datasets of increasing size shows that, although small datasets of 25 plates may miss some modes, larger datasets allow REPOP to accurately recover the underlying multimodal structure in both cases. We quantify this behavior further in Appendix 2, where we analyze the error between the reconstructed and ground-truth means, as well as the relative entropy between the reconstructed and ground-truth distributions, as a function of the number of plates. Together, these results demonstrate REPOP’s ability to infer the true population despite stochastic noise introduced by dilution and plating.

In Fig. 3a, we show a relatively simple dataset, where the three peaks are already discernible in the histogram obtained via naive multiplication. Nonetheless, REPOP improves the resolution of these peaks, particularly as the number of samples increases. In Fig. 3b, we present a more challenging case where the modes are less distinct under the naive multiplication approach; the additional stochasticity introduced by dilution causes the resulting histogram to appear as a single broad peak. Nonetheless, REPOP is able to accurately recover the underlying multimodality. In both cases, the performance of REPOP improves with larger datasets, as evidenced by the progressively sharper resolution of each peak. We quantify this improvement in Appendix 2, where we analyze the reconstruction error in the mixture means and the Kullback-Leibler (KL) divergence between the reconstructed and ground-truth distributions as functions of the number of plates.

### When the cutoff is taken into account

For our final demonstration with simulated data, we show results when the data follows a cutoff in reporting, as described in the likelihood for Model 3. We have generated simulated data according to the following scheme: starting from the lowest dilution (20, as shown in Fig.1a). If the counts are larger than the cutoff, we used *k*_CO_, we sample again at a dilution factor 10 times higher until we find a plate that has fewer colonies than *k*_CO_. We then record that number of counts and that dilution factor as the *i*-th datapoint (*k*_*i*_, *ϕ*_*i*_), as would be done by rigorously following the cutoff. If we know the dilution schedule, this is equivalent to having the data as the counts and the index of the reported dilution (*k*_*i*_, *m*_*i*_) as in (17), with each reported dilution matched to the corresponding index *m*_*i*_, meaning the dilution reported is the *m*_*i*_-th in the schedule 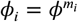.

We visualize the data generated this way and show the reconstruction obtained for Model 3 in Fig. 4. In the rightmost column, we demonstrate how failing to account for the cutoff leads to an artificial peak (*e*.*g*., an extra mode near the leftmost peak in the ground truth, Fig. 4b). This occurs because counts near the cutoff *k*_CO_ are ambiguously assigned to incorrect dilutions.

**Figure 4.**
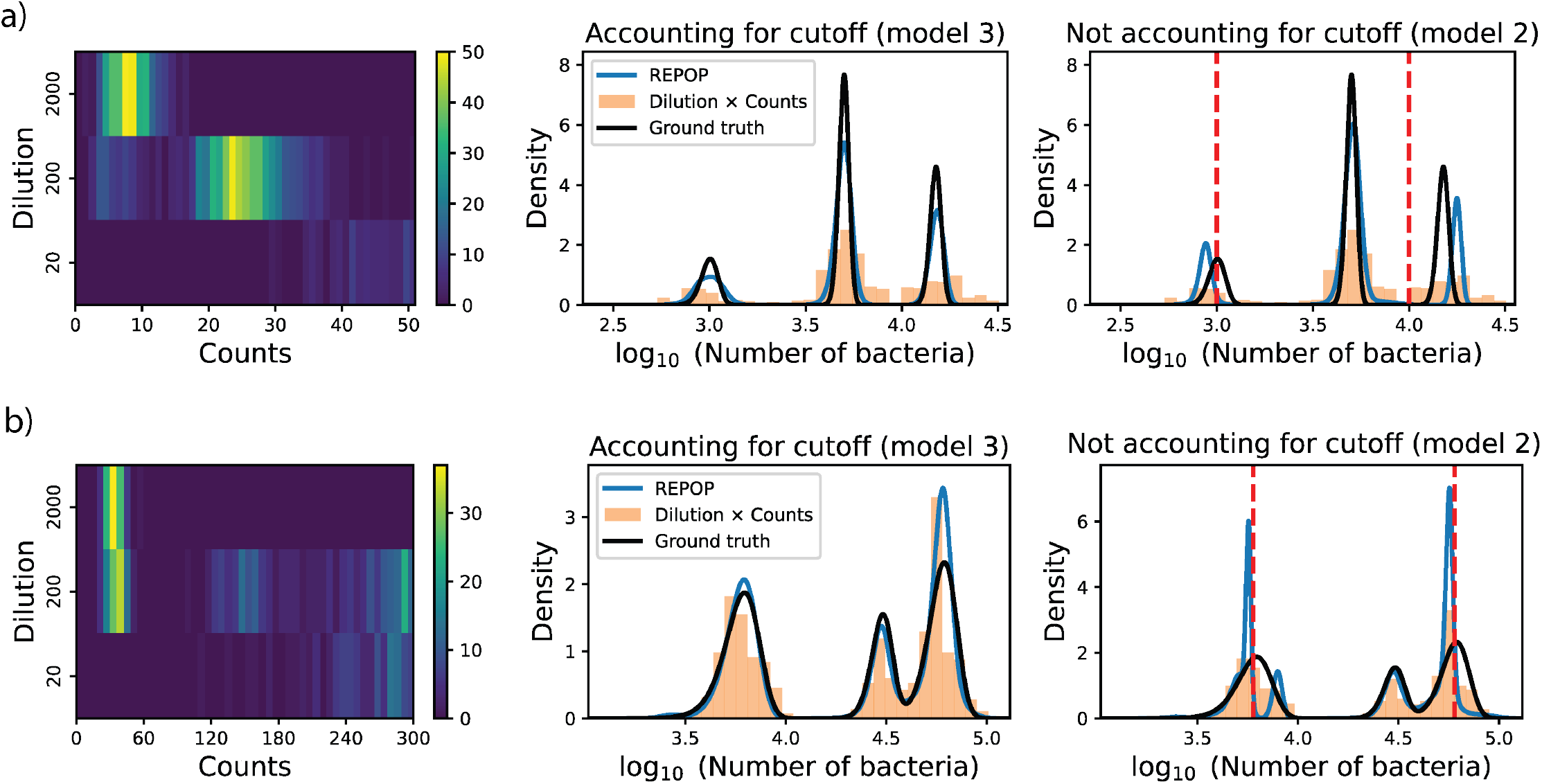
Not accounting for the cutoff can lead to incorrectly attributed multimodality. This figure compares population reconstructions with and without accounting for the cutoff effect, (using Model 3 and Model 2 respectively). a) simulated data generated with a cutoff *k*_CO_ = 50. The ground truth population consists of three Gaussian components with means 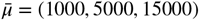, standard deviations 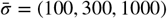, and weights 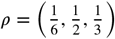 A more realistic cutoff *k*_CO_ = 300, with ground truth parameters 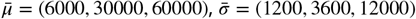, and 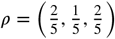 . The middle column shows reconstructions using Model 3 (which accounts for the cutoff), while the rightmost column shows results from Model 2 (which ignores it). The rightmost column also highlights key cutoff crossings, using red dashed lines. These lines indicate where bacterial counts approach the cutoff at different dilutions. Notably, failing to account for the cutoff leads to incorrect mode placement, including the emergence of an extra peak observed in b). As discussed in the main text, when the true distribution has substantial probability near these points (meaning there is a non-negligible probability to find a sample with these number of bacteria close to those cutoff values) failing to account for the cutoff effects results in inaccurate reconstructions.

These errors become particularly pronounced when colony counts approach *k*_CO_, as the cutoff procedure forces certain samples to be reported at different dilutions. For example, consider a cutoff *k*_CO_ = 300 and two possible dilution factors, *ϕ*^1^ = 20 and *ϕ*^2^ = 200. A sample with an original bacterial count *n*_*i*_ near *k*_CO_*ϕ*^1^ = 6000 may yield counts at dilution *ϕ*^1^ that either exceed or fall below the cutoff due to variability in the plating process. When the cutoff effect is ignored, the algorithm mistakenly interprets these counts as coming from separate populations, leading to incorrect peak formation, as seen in the rightmost column of Fig. 4b. Beyond this qualitative observation, we also assess the magnitude of these errors by computing the relative deviation between the estimated and ground truth means for each component (see Appendix 2).

### Results for experimental data: Discerning different true concentrations of bacteria

While we have demonstrated in the previous subsections how to resolve overlapping distributions in simulated datasets, here we move on to obtaining a similar case built with real-world plate counting. In particular, we construct a case that emulates multiple peaks at different dilutions, as shown in Fig. 4, by generating four different sample sources of bacteria.

We prepared vials with four different concentrations of *E*.*coli* measured by optical density, and consider samples of a volume equivalent to 0.2 nL from these vials. These samples are further diluted according to the dilution schedule Φ = 20, 200, 2000. However, in practice, we do not directly dilute 0.2 nL. Instead, we take 1 *µ*L and dilute it by factors of 10^5^, 10^6^, and 10^7^ – further details in Appendix 3. This approach ensures that while the original vials can be assumed to have negligible variability, the small 0.2 nL subsamples may no longer be, resulting in a unimodal but with a larger relative variance population for each vial.

The colony counts obtained are processed without identifying their original vial, aiming to reconstruct the underlying four components structure. However, the dataset remains relatively small, consisting of only 97 total samples (see details in Appendix 3). This dataset and the reconstructed distribution are presented in Fig. 5a.

**Figure 5.**
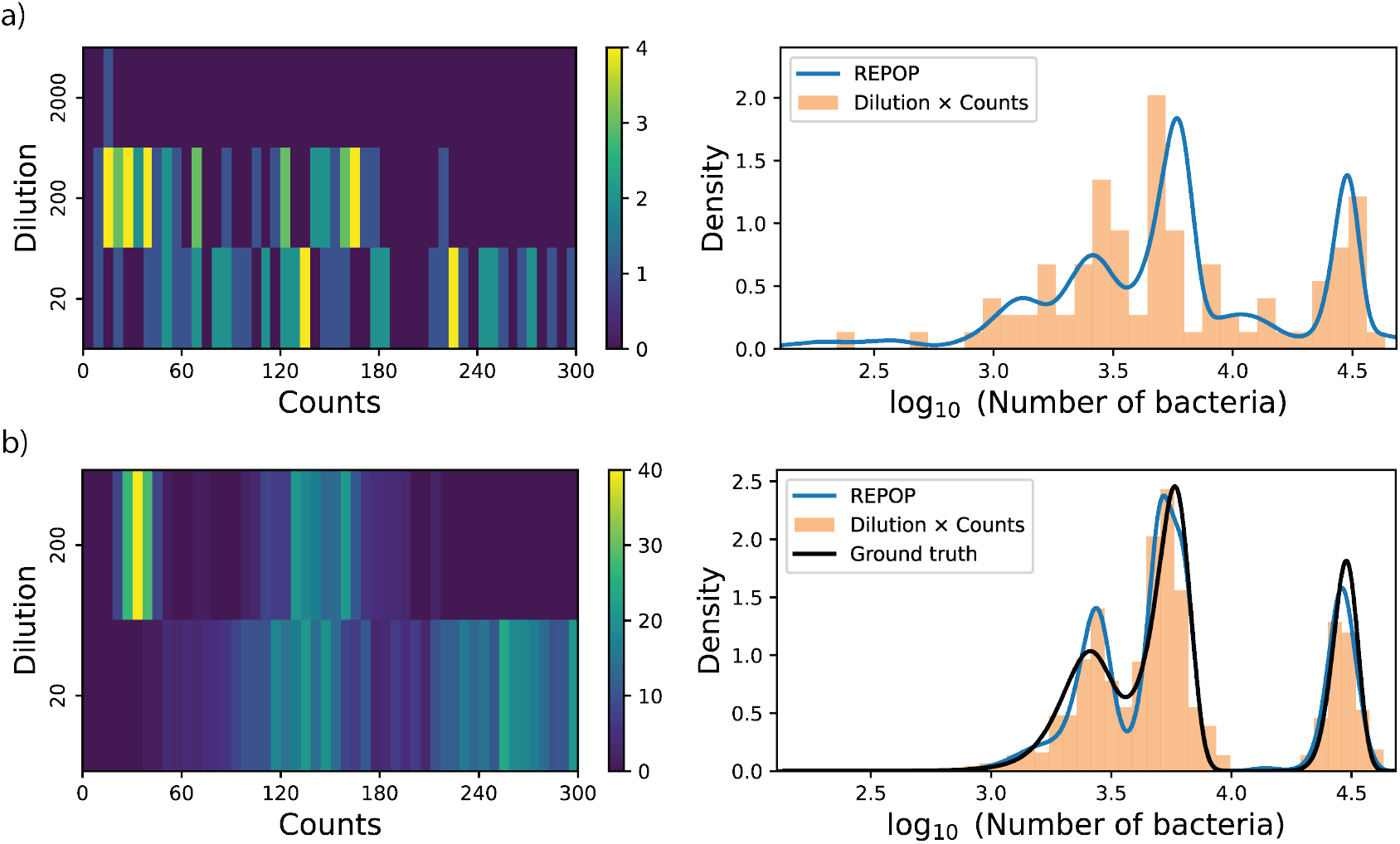
Reconstructing, from real experimental plate counting, the population from vials with different optical densities. a) Distribution of observed colony counts obtained as described in text. The data is passed to the system without vial identification, aiming to reconstruct the underlying structure. b) Results from a simulated dataset with 750 datapoints (plates). The increased sample size improves the estimation of the four main components, (which appear as approximately three peaks because two components substantially overlap), yielding a more accurate reconstruction.

Although the limited sample size results in an imperfect match, the method successfully captures the four main components – corresponding to the estimated values of *µ*_*α*_ and *σ*_*α*_ associated with the four highest values of *ρ*_*α*_. To further validate the approach, we supplement the dataset by generating 750 synthetic datapoints from the mixture distribution made of the four main components, leading to a dataset considerably larger than what is typical of plate counting. The results, shown in Fig. 5b, demonstrate a significant improvement in capturing the underlying structure of the data.

### Application: Capturing host differences through examining gut microbiota population

Here, we use REPOP to studying heterogeneity in more complex biological systems, such as microbial ecology within a host organism. Thus, we apply our framework to analyze gut bacterial counts from individual *C. elegans*.

Heterogeneity in bacterial counts among individual *C. elegans* has been reported and is typically attributed to rare colonization events and size variability (***Vega and Gore, 2017; Taylor and Vega, 2024; Eisenmann, 2005***). However, to derive meaningful insights into gut ecology, it is essential to distinguish true population heterogeneity from the stochasticity introduced by dilution and plating.

To minimize confounding factors, we restrict our analysis to the OP50 strain of *E. coli*, which is well-characterized in laboratory settings and, under idealized conditions, is expected to produce a unimodal population distribution. Any deviations from this expected pattern may reflect individual *C. elegans* variability, either due to differences in the gut microenvironment or the fact that gut colonization is a rare event (***Vega and Gore, 2017***).

Due to this, we conducted an experiment in which *C. elegans* were synchronized, cleared of gut bacteria at three days post-hatching and subsequently fed live *E. coli* for controlled durations (1, 3, 5, 7, or 9 days). At each studied time point, 25 individual worms were picked, cleaned of all non-gut adhered bacteria, and crushed. The worm homogenate was then serially diluted and plated with a dilution schedule, Φ = {22, 222, 2222}. The plates were counted after a 24 hour incubation. More detailed experimental methods may be found in the Appendix 3.

The results presented in Fig. 6 demonstrate a clear trend in bacterial population dynamics within the *C. elegans* gut. The population distribution shifts consistently to higher values with each successive day, aligning with our expectation that bacterial numbers increase over time due to colonization and replication inside the gut.

**Figure 6.**
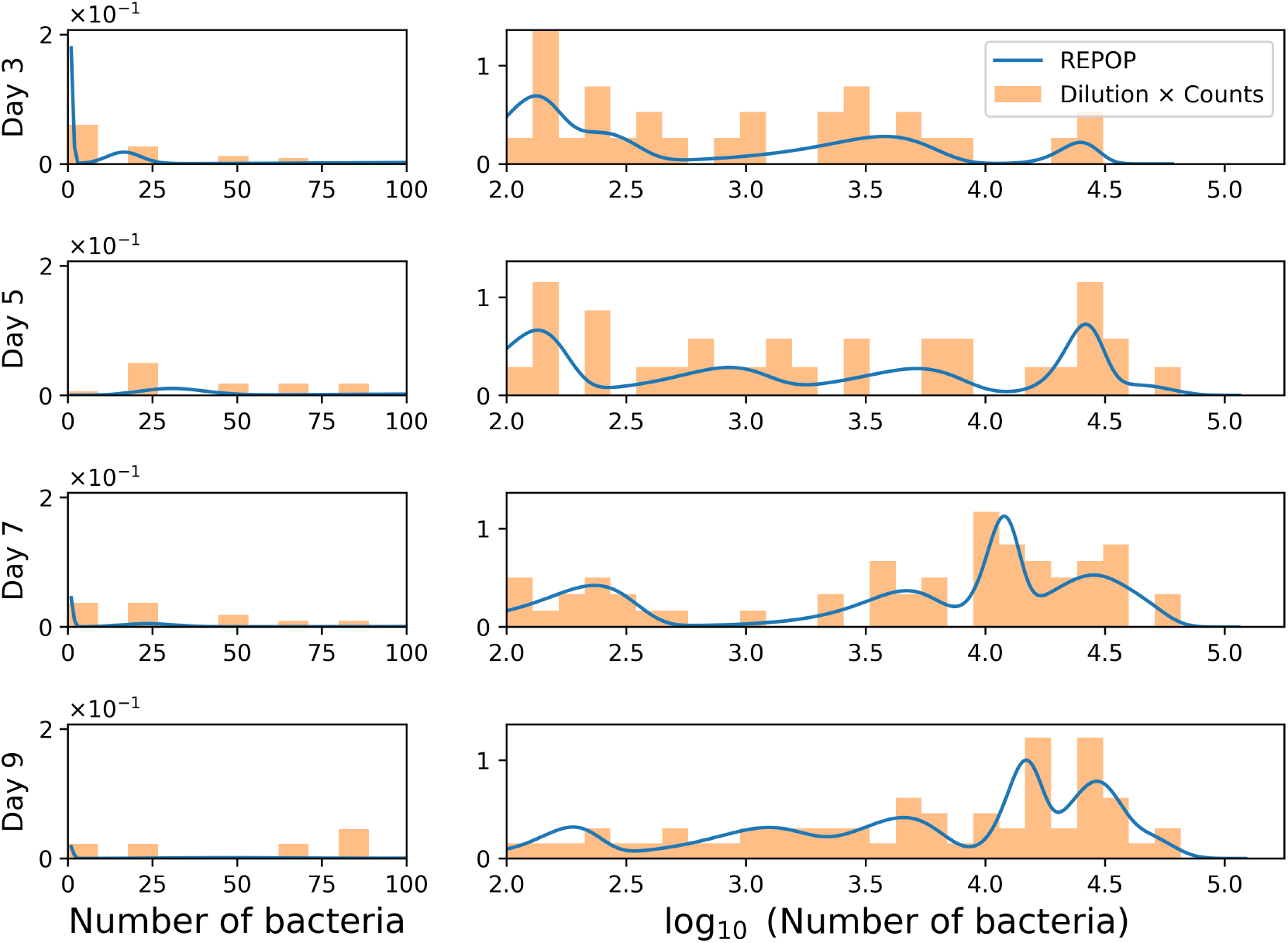
Results for the population of *E.coli*-fed *C. elegans*’ gut at days 3, 5, 7, and 9 of adulthood. Here we are able to observe the shift in the population distribution obtained with plate counting shifts to higher population for longer-living *C. elegans*. To better visualize the distribution and model fits, we use a split-log x-axis: bacterial counts from 0 to 100 are shown on a linear scale, and counts above 100 on a logarithmic scale. The population distribution shifts toward higher values each day, as expected from colonization and replication within the gut. If biological significance, such as different *C. elegans* types, as proposed in *e.g*. (***Boddu et al., 2024***) is to be inferred from this multimodality, a rigorous separation of population heterogeneity and plating stochasticity, as implemented by REPOP, is essential.

The multimodality exhibited in Fig. 6 can be explained by the heterogeneity of the host. Recently, this has been corroborated by *in vivo* experiments (***Boddu et al., 2024***). This phenomenon could be related to the well-established stratification in heat-shock protein (HSP) expression, which is also correlated with aging and lifespan in nematodes (***Yashin et al., 2002; Wu et al., 2006; Vertti-Quintero et al., 2021***). Thus the variation in the population could be due to the fact that ingestion and egestion exhibit an age-related change (***Bolanowski et al., 1981; Huang et al., 2004***) or due to state switching (***Boddu et al., 2024***) in feeding and digestion, akin to genetic state switching (***Pessoa et al., 2024; Kilic et al., 2023***). The results in Fig. 6 support the conclusion that biologically meaningful differences in population counts are being captured, rather than the data being dominated by instrumentation error. REPOP provides a robust method for distinguishing multiple peaks in the population distribution, indicating that intrinsic variability among individual *C. elegans* can be deconvolved from measurement noise.

### Resolving different species

One possible extension of plate counting is the quantification of multiple bacterial species or phenotypes within the same biological sample. In practice, this could be done when different bacteria produce distinguishable colony colors or morphologies (***Maeda et al., 2017; Qamer et al., 2003; Haldeman and Amy, 1993***). In such cases, the counts of different bacterial species can be obtained separately from the same sample.

To illustrate how the framework can be extended to this setting, we consider a synthetic dataset consisting of two correlated species. For each sample *i*, we denote by 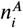 and 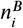 the unobserved numbers of bacteria of species *A* and *B*, respectively, drawn from a joint distribution that captures their correlation. Correspondingly, the plate count measurements yield observations 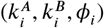. In this section, we use these synthetic data to examine how well the framework can recover the marginal population distributions of the two species from their respective plate count measurements.

In the example shown in Fig. 7, we generate a synthetic two-species population distribution with three modes. The mixture components have equal weights and means (4000, 24000), (14000, 8000), (8000, 16000), where the first and second coordinates denote the population sizes of species *A* and *B*, respectively. The covariance matrices are chosen so that the marginal distribution of species *A* matches the trimodal distribution used in Fig. 1c, while the marginal distribution of species *B* consists of three approximately equally spaced modes at 8000, 16000, and 24000, with equal variances. In the ground truth distribution, two of the mixture components include correlations between the populations of species *A* and *B*, as seen in the population samples shown by the black points in Fig. 7.

**Figure 7.**
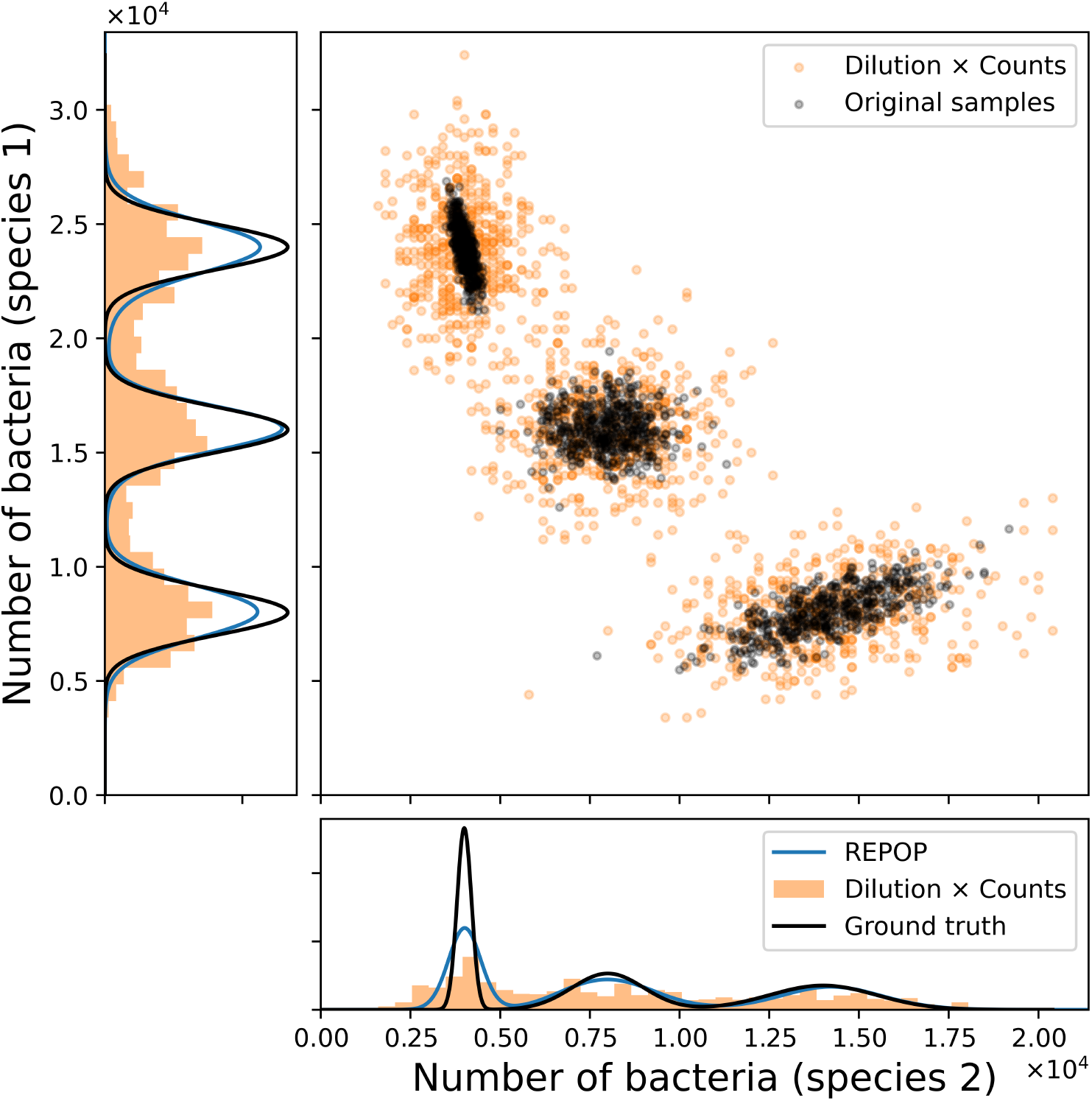
REPOP reconstructs marginal population distributions when individual plates distinguish two species. A synthetic dataset of 1500 samples contains two correlated bacterial populations. Black points denote the true underlying populations (*n*^*A*^, *n*^*B*^ ), which are drawn from a joint distribution with three well-separated modes, as described in the text. Orange points denote the corresponding plate counts after a dilution factor of 200, naively reconstructed as counts multiplied by the dilution factor. Although the true population (black) modes are well separated, stochasticity introduced by dilution and plating causes the observed measurements(orange) to overlap. We assume that the two species can be distinguished on each plate, so that separate counts are obtained for species *A* and *B*. Applying REPOP separately to the counts from each species recovers the corresponding marginal population distributions, as shown in the side panels.

We then simulate plate count measurements from this population distribution using a dilution factor *ϕ*_*i*_ = 200, generating a synthetic dataset of 1500 samples. As shown in Fig. 7, the true population modes are well separated. However, after dilution and plating, the naively reconstructed observations, obtained by multiplying the plate counts by the dilution factor, are broadened and partially overlap. Nevertheless, applying REPOP separately to the species-specific counts recovers the marginal population distributions for both species.

## Conclusion

In accurately estimating bacterial counts, simply multiplying counts by the dilution factor provides only a rough first approximation. As we showed here, this method alone often fails to capture the original individual counts across samples, as straightforward multiplication of colony counts by the total dilution factor typically leads to an overestimation of the spread (Fig. 1b) that may obscure multimodal behavior (Fig. 1c).

This is not merely a technical issue in quantification: it directly affects the biological interpretation of the data. For example, inferring population growth dynamics (***Armitage and Jones, 2019; Martino et al., 2016; Taylor and Vega, 2024; Martini et al., 2024; Boddu et al., 2024***) and learning features driving population heterogeneity across samples (***Armitage and Jones, 2019; Goberna and Verdú, 2022; Pinto et al., 2022***), demands a deeper analysis of the statistics involved in dilution and counting. Indeed, such a treatment avoids attributing ecological (or other) significance to the randomness introduced by plating.

The Bayesian reconstruction presented here and implemented through REPOP provides a rigorous method to obtain the distribution of populations from plate counts, including observing the correct multimodality of the population, as seen in Figs. 1 – 4. This allows the method to extract more information from each individual plate and to reconstruct population features that may be obscured in the raw observations, including multimodality and distributional differences across conditions. This setting is naturally suited to a Bayesian treatment because the quantity of interest is not the observed colony count itself, but the latent population distribution that gave rise to the observed colony counts.

The study of the multimodal case accounting for cutoff (Model 3) also demonstrates that retaining all data, regardless of their individual importance in reconstructing the final population size distribution, remains crucial. As shown in Fig. 4, ignoring the cutoff can lead to misidentifying multimodality, especially for plates with colony counts near the cutoff. Consequently, REPOP’s results emphasize that explicitly reporting both the dilution schedule and the cutoff values used should be a fundamental part of data reporting, as they impact the form in which likelihoods are calculated.

Moreover, due to REPOP’s joint statistical description of dilution and plating, REPOP can be extended beyond the specific cases considered here. In particular, the same treatment of measurement stochasticity can be combined with alternative population models designed for more specific ecological hypotheses, as explored in related work by some of us (***Lu et al., 2026***). In that work, the joint measurement model was combined with a specific parameterized ecological model, allowing posterior inference over *p*(*θ* ∣ *D*) and therefore uncertainty quantification over the model parameters. Specifically, the REPOP measurement model was used to provide the measurement probabilities *p*(*k*_*i*_, *ϕ*_*i*_ ∣ *n*_*i*_), as described in the section “Population versus measurement stochasticity.”

Therefore, while the main implementation of REPOP focuses on finding the population distribution that maximizes the posterior, as described in (18), and is designed to remain model-agnostic by using Gaussian or mixtures of Gaussians as flexible models for the population distribution, its measurement model can also be incorporated into hypothesis-specific population models. This makes REPOP not only a practical reconstruction tool but, more importantly, a foundation for future inference problems involving plate counting data.

Furthermore, to help with computational scalability, REPOP can optionally as specified by user rely on GPU acceleration. This can become helpful as likelihood expressions in (8) and (9) require summations over all possible values of the underlying population size, which can become memory and time intensive. REPOP mitigates these computational demands through GPU parallelization, as performance can be improved substantially from CPU-only systems. Moreover, extending the framework to jointly infer multiple interacting populations in the future would increase computational cost considerably, since the relevant sums must then be performed over the product space of possible population values. In the species case, this leads to an effective quadratic scaling with population range. This quickly becomes prohibitive for larger populations, which is why, in Fig. 7, we focus on marginal rather than full joint distributions. Developing more efficient strategies for multiple species inference problems remains an important direction for future work.

Beyond that, a natural extension of the present framework is to refine the measurement model itself by incorporating additional sources of counting error beyond dilution and plating stochasticity. For instance, integrating the notion of additional error sources, such as false positive or negative rates in counting that due to *e.g*. bacterial colony overlap, could help in further reducing the breadth in the original counts over bacteria. For instance, intuitively, we may imagine the probability of miscounting two or more colonies as a single one grows as we reach colony counts near the cutoff. Simple simulations, for instance, considering typical colony and plate size, may be used to gain initial insight on potential false negative rates, and thus its effect on the original sample size.

Finally, REPOP provides a means by which to test hypotheses (***Boddu et al., 2024; Martini et al., 2024***) relating observed multimodality in gut microbial content of *C. elegans*, as shown in Fig. 6, to distinct categories of hosts. What is more, REPOP can also be used to study other organisms, exhibiting similar multimodality in their microbiota (***Acuña Hidalgo and Armitage, 2022; Clemmons et al., 2015; Chen et al., 2024***). The central point is that identifying biologically meaningful heterogeneity requires more than collecting large datasets: it requires a framework that distinguishes true ecological structure from variability introduced by the measurement itself. Without such a framework, one risks assigning ecological significance to artifacts of dilution and plating. REPOP is designed precisely to avoid those biases while remaining flexible enough to accommodate complex and multimodal population structure.

## Acknowledgments

SP acknowledges support from the NIH (R35GM148237), ARO (W911NF-23-1-0304), and NSF (Grant No. 2310610). Figures 1a and 2 were created in BioRender. Presse, S. (https://BioRender.com/espzw5y) is licensed under CC BY 4.0. We also thank Ayush Saurabh, Max Schweiger, Lance (W. Q.) Xu, and Ioannis Sgouralis for insightful discussions in the development of this work.

## Appendix 1

### Prior selection across the unimodal and non-parametric models

When estimating the non-parametric model by MAP, we use a prior *p*(θ) designed to accommodate multimodal population distributions. Here, we describe this prior choice in detail.

As mentioned in the main text, this prior encompasses all of the parameters needed to construct the probability distribution for *n*, given by Eq.(5) in the main text. Therefore, θ must include the means 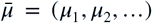, standard deviations 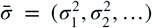, and mixture weights 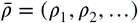of the Gaussian mixture, 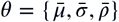. As mentioned in the main text, for practical purposes we assume a weak limit, which we refer to as *A* and equivalent to the truncation on the number of components in Eq. (6) in the main text (defaulted as the smallest value among 25 and the square root of the total number of samples) although it is changeable in our software.

Let us separate each piece of the prior 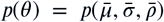. First, we consider the prior for the weights. In a uni-modal distribution, case 1, the result is trivial: we have ρ_1_ = 1 with probability 1. However, in a multi-modal cases (2-3) where the number of components is unknown, we use a Dirichlet distribution, a common choice for mixture models in Bayesian non-parametrics ***Pressé and Sgouralis*** (***2023***), given by:

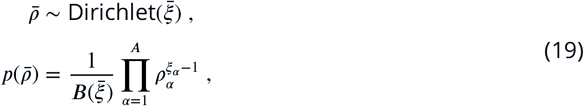

Where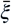is a sequence of the same shape as 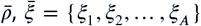 and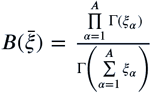 with Γ representing the gamma function. In order to prevent overfitting is an usual choice to have an exponentially decreasing. Let us write 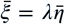 with 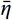 being a sequence 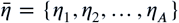 where the parameters decay exponentially by a factor *q*, such that η_α_ = *q*^α^, and λ being a scalar (we use 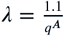 ). The expected value of *ρ*_*α*_ sampled from (19) will be 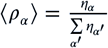.

With the prior for 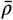 established, we proceed to define the prior for the means 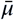 and standard deviations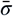 . Here, each component *α* is independent from each other. Initially, for the means, we select a distribution that avoids biasing the choice by being nearly uniform, but preventing too small numbers and adhering to a sensible cutoff in *n*. The mean of each Gaussian must be smaller than twice the largest number found by naively multiplying counts per dilution; we denote this largest cutoff as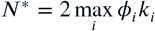. Thus, the prior for each mean, *µ*_*α*_, is given by

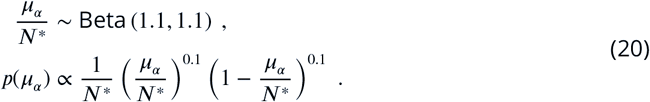

For the prior in 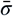, we write a prior that scales with its respective means, but is still broad; we choose a lognormal prior here

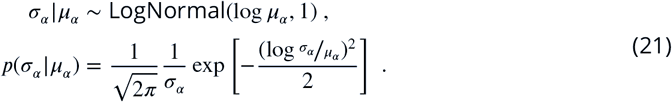

Thus, the prior used to generate the figures in the main text is defined as:

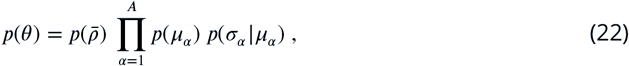

with 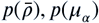, and *p*(*σ*_*α*_|*μ*_*α*_) given by (19), (20), and (21) respectively.

## Appendix 2

### Error metrics between reconstructed and ground-truth distributions

In Figs. 3 and 4, we showed that REPOP can reconstruct underlying population distributions obscured by dilution and plating stochasticity. Here, we make this comparison quantitative using two complementary error metrics. The first measures how well the inferred mixture components recover the locations of the ground-truth modes. The second compares the full reconstructed and ground-truth distributions using the KL divergence.

To compare two distributions, *p*(*n*_*i*_|*θ*) and *p*(*n*_*i*_ *|θ_GT_*), in terms of how well their component means match, we use the relative error between the means defined as

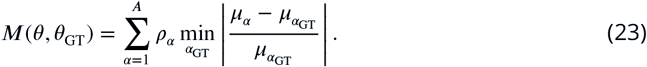

Here, *p*(*n_i_ |θ*) is the reconstructed distribution (through REPOP) and *p*(*n*|*θ*_GT_ ) is the groundtruth distribution. The index *α* labels the components of the reconstructed distribution, while *α*_GT_ labels the components of the ground-truth distribution. The quantity inside the absolute value is the relative error between the reconstructed component mean *µ*_*α*_ and a ground-truth component mean 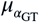 . We take the minimum over α_GT_ to account for cases in which component labels do not match across the two distributions. The weights ρ_α_ are the reconstructed mixture weights, so components with larger inferred probability mass contribute more strongly to the metric.

To compare the full reconstructed and ground-truth distributions, rather than only the locations of their modes, we use a standard metric for comparing probability distributions: the relative entropy, also known as the Kullback-Leibler (KL) divergence. For each reconstructed distribution *p*(*n*| *θ*) and corresponding ground truth distribution *p*(*n* |*θ*_GT_), we define

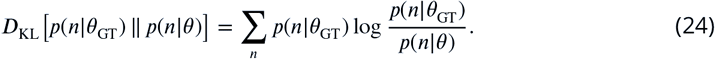

The sum is taken over the population size *n*. Here, *p*(*n* | *θ*_GT_) is the known ground truth mixture distribution used to generate the simulated data, while *p*(*n*|*θ*) is the mixture distribution reconstructed by REPOP. Roughly, KL divergence measures how much information is lost when the reconstructed distribution is used as an approximation to the ground truth distribution. In the figure below, we show how both metrics decay as the dataset size increases for the same examples shown in Fig. 3.

Analogously, for the dataset shown in Fig. 4a, the relative error *M* in (23) is 1.50% when the cutoff in the total number of colonies is accounted for (model 3). When the cutoff is ignored (model 2), the error increases substantially to 7.47%. The same trend is observed for the KL divergence in (24), which increases from 0.13 when the cutoff is included to 1.73 when the cutoff is ignored. Similarly, for the dataset shown in Fig. 4b, the relative error is 2.07% when the cutoff is considered, compared with 11.5% when it is not. The KL divergence likewise increases from 0.03 to 0.64 when the cutoff is ignored. Together, these results show that accounting for the cutoff is necessary to accurately reconstruct both the mode locations and the full population distribution. Full details of these calculations are available in our GitHub repository ***Pessoa*** (***2025***).

**Appendix 2—figure 1.**
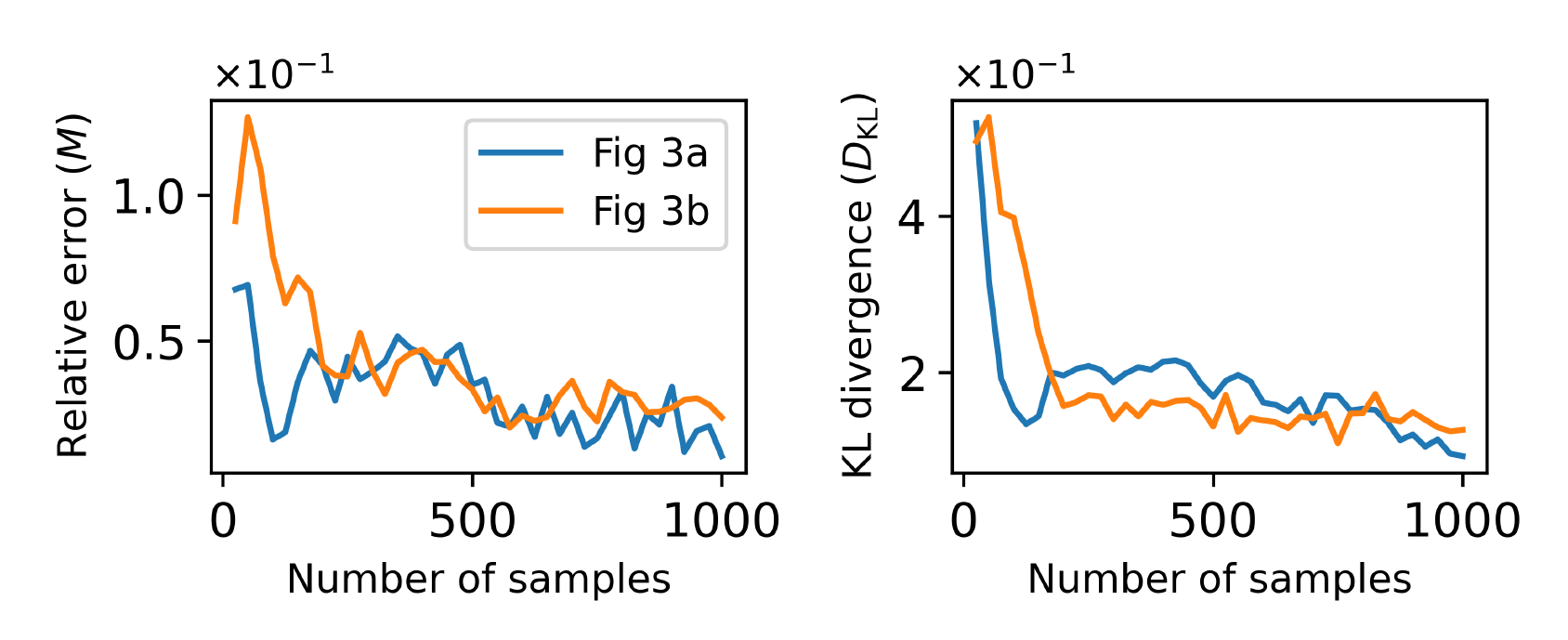
Decay of error metrics as the dataset size increases for the distributions shown in Fig. 3. We show the relative error defined in (23) (left) and the KL divergence defined in (24) (right) for the distributions in Fig. 3a and Fig. 3b. Both metrics decrease as the number of plates increases, indicating improved agreement between the REPOP reconstruction and the ground-truth.

## Appendix 3

### *E. coli* culture for simulating different expected population peaks

An overnight culture of the OP50 strain of *Escherichia coli* was grown in lysogeny broth (LB) media, and its optical density at 600 nm (OD_600_) was measured against an LB blank. The culture was concentrated to an expected 5X of OD_600_=1.0 by centrifugation and resuspension in phosphate-buffered saline (PBS). The 5X suspension was then diluted to an expected OD_600_ of 0.5, 0.1,, and two at 0.01, with empirical OD_600_ values recorded against a PBS blank, which were 0.804, 0.188., 0.025, and 0.02. From the first three vials, we obtained 25 samples (each with 3 dilutions), while the last vial yielded 22 samples.

For each OD_600_ condition, serial dilutions were prepared to 1:100,000, 1:1,000,000, and 1:10,000,000 in PBS. A total of 90 *µ*L from each dilution was plated onto LB agar, resulting in 75 plates per OD condition. Plates were incubated at 37°C for 18–24 hours before counting colony-forming units (CFUs) to estimate bacterial concentration.

### *C. elegans* culture

The temperature-sensitive, reproductively sterile strain AU37 of *Caenorhabditis elegans* (*C. elegans*) was cultivated and synchronized according to standard protocols (***Eisenmann, 2005***). Following the protocol described by Gore and Vega (***Vega and Gore, 2017***), synchronized eggs were cultivated at the non-permissive temperature of 25°C to produce sterile adult worms. These worms were then washed and transferred to a heat-killed E. coli OP50 suspension in S media supplemented with gentamicin for 24 hours to clear gut bacteria.

Once cleared of gut bacteria, worms were washed once with 10 mL M9 buffer containing 0.1% Triton X and twice with 10 mL M9 buffer. They were then transferred to a live bacterial suspension (E. coli OP50 at OD_600_ = 1.0 in 1 mL S media) for a duration of 3, 5, 7, or 9 days. Fresh bacterial suspension was provided at 24, 72, 120, and 168 hours (approximately every 48 hours after the initial 24-hour feeding).

On the designated sampling days for gut bacterial enumeration, worms were washed to remove non-adhered surface bacteria and manually disrupted following the protocol outlined in the section titled *Disruption of Worms and Plating of Intestinal Populations* of ***Vega and Gore*** (***2017***). The resulting homogenate was serially diluted using a 10-fold dilution series, with final expected dilution factors of 1:22, 1:222, and 1:2222 before plating. Colony-forming units were enumerated after incubating the plates for 24 hours.

## Notes

https://github.com/PessoaP/REPOP

### Competing Interest Statement

The authors have declared no competing interest.

### Summary of Updates

In response to a series of reviewer on eLife we added: - A new case study showing a framework application to two interacting species. - A clearer distinction between measurement stochasticity and ecological variability, together with a discussion of how the framework can support future ecological inference problems. - Additional clarification of the software's use, implementation, and underlying assumptions.

